# Estradiol Signaling in Regulatory T Cells Coordinates the Remodeling of the Maternal Small Intestine during Pregnancy

**DOI:** 10.64898/2026.11.11.765966

**Authors:** Luting Liu, Xianyang Jin, Jiaman Zhang, Shangbo Tian, Yujie Wang, Liuyong Lu, Xuemei Jiang, Yong Zhuo, Chao Jin, Bin Feng, Lianqiang Che, Shengyu Xu, Yan Lin, Long Jin, Mingzhou Li, De Wu, Lun Hua

## Abstract

Pregnancy requires coordinated maternal adaptations to support fetal growth, yet signals that couple gestation to intestinal remodeling remain poorly defined. Here, we show that pregnancy induces a reversible regenerative program in the small intestine characterized by villus elongation, increased nutrient transporter expression, and enhanced absorptive capacity driven by intestinal stem cell (ISC) hyperproliferation. Using single-cell transcriptomic and genetic approaches, we identify intestinal regulatory T (Treg) cells as required regulators of this remodeling. Estradiol treatment of ovariectomized mice fully recapitulated the gestational intestinal remodeling phenotype, whereas Treg-specific deletion of estrogen receptor α reduced villus expansion and impaired nutrient absorption. Mechanistically, elevated estradiol levels during pregnancy expand Treg cells and enhance IL-10 production, which directly promotes ISC proliferation and epithelial renewal. Cross-species analyses in mice and pigs support conservation of the intestinal phenotype, while integration of human GWAS data links Treg-associated IL-10 signaling to birth-weight-associated traits. Together, these findings demonstrate that pregnancy repurposes immune tolerance pathways to coordinate maternal intestinal remodeling with fetal nutritional demands.

## Introduction

Pregnancy imposes substantial physiological demands that require coordinated maternal adaptation to support fetal growth and development. To meet these demands, multiple maternal organs undergo extensive functional and metabolic remodeling, including the liver, adipose tissue, pancreas, cardiovascular system, and immune system. Together, these adaptations reprogram maternal metabolism to maximize nutrient acquisition, storage, and transfer to the developing fetus (***Falick Michaeli et al., 2015; Mor, Aldo, & Alvero, 2017; Sanghavi & Rutherford, 2014; Trowsdale & Betz, 2006; Yang et al., 2025; Yu et al., 2024***) The gastrointestinal tract, particularly the small intestine, also undergoes structural and functional remodeling that markedly enhances nutrient absorption (***Ameku et al., 2025; Huang et al., 2026; Onji et al., 2025***) However, signals that couple gestation to intestinal remodeling remain largely unknown.

The intestinal epithelium is one of the most rapidly self-renewing tissues in adult mammals. This continuous renewal depends on the proliferation and differentiation of intestinal stem cells (ISCs) residing within the crypt niche (***Santos, Lo, Mah, & Kuo, 2018; Shay & Yilmaz Ö, 2025; van der Flier & Clevers, 2009***). ISC activity responds to epithelial and mesenchymal niche cells and also to resident immune cells that act on epithelial regeneration and tissue homeostasis (***Biton et al., 2018; Chi et al., 2025; Gronke et al., 2019; Lindemans et al., 2015; Sullivan et al., 2021***). Pregnancy also induces immunological adaptations that establish maternal–fetal tolerance while maintaining protective immunity (***Aghaeepour et al., 2017; Apps et al., 2020; Chen et al., 2022***). Regulatory T (Treg) cells undergo systemic expansion during pregnancy and are essential for maintaining maternal–fetal immune tolerance (***Aluvihare, Kallikourdis, & Betz, 2004; Li et al., 2025; Rowe, Ertelt, Xin, & Way, 2012; Samstein, Josefowicz, Arvey, Treuting, & Rudensky, 2012; Somerset, Zheng, Kilby, Sansom, & Drayson, 2004***). However, whether these pregnancy-induced immune adaptations also contribute to the physiological remodeling of peripheral metabolic organs, including the intestine, remains largely unexplored.

Pregnancy also induces endocrine remodeling. Circulating levels of reproductive hormones, including estradiol, progesterone, and placental hormones, rise progressively throughout gestation, particularly during late pregnancy, and contribute widespread maternal physiological adaptations (***Ammari et al., 2023; Soldin et al., 2005; Tan & Tan, 2013; Wu, Li, & DeMayo, 2018***). Recent studies suggest that these hormonal changes promote regenerative responses in the maternal intestine (***Ameku et al., 2025; Beumer & Clevers, 2016; L. Liu et al., 2023; Onji et al., 2025***), however, the specific hormones, target cell populations, and underlying molecular mechanisms that mediate this process during pregnancy remain poorly defined. Given that endocrine and immune adaptations occur in parallel throughout gestation, hormonal signals may reprogram immune cell function to coordinate tissue-specific regeneration. However, whether endocrine signaling acts directly on immune cells to drive ISC-dependent intestinal remodeling remains unknown.

Here, we identify an endocrine–immune signaling axis that links maternal hormonal adaptation to intestinal remodeling during pregnancy. Using single-cell transcriptomics, genetic mouse models, intestinal organoids, and functional analyses, we demonstrate that gestational estradiol signaling acts through estrogen receptor α (ERα) in intestinal Treg cells to enhance IL-10 production, thereby promoting ISC proliferation, epithelial renewal, and nutrient absorption. Integrative analyses of human genome-wide association studies further reveal associations between Treg-associated IL-10 signaling pathways with birth-weight-related and pregnancy-associated metabolic traits, supporting the physiological relevance and cross-species conservation of this pathway. Together, our findings define an estradiol–ERα–Treg–IL-10–ISC signaling axis that integrates maternal endocrine and immune adaptations to coordinate intestinal remodeling during pregnancy and identify immune-mediated epithelial remodeling as a maternal adaptation that supports fetal growth.

## Results

### Pregnancy induces conserved intestinal remodeling across the mammalian species examined to enhance maternal nutrient absorption

To investigate how the maternal intestine adapts to the increasing metabolic demands of pregnancy, we examined intestinal morphology, epithelial architecture, and absorptive function in age-matched virgin female mice, pregnant mice at embryonic day 18.5 (E18.5) of gestation, and dams after a 14-day recovery period following weaning (21 days of lactation followed by pup removal). This experimental design allowed us to distinguish pregnancy-specific intestinal adaptations from persistent physiological changes following pregnancy.

Consistent with previous reports (***Ameku et al., 2025***), pregnancy significantly increased both the length and mass of the small intestine compared with virgin mice (***Figure 1-supplementary figure 1A and B***). Histological analysis further revealed marked villus remodeling during late gestation, with significant increases in both villus height and villus width (***Figure 1A and B*; *Figure 1-supplementary figure 1C***). These structural adaptations largely regressed after weaning recovery, although several morphological parameters remained modestly elevated compared with virgin mice, indicating that pregnancy-induced intestinal remodeling is largely, but not entirely, reversible. Quantitative morphometric analysis further showed that pregnancy significantly increased epithelial cell size throughout the villus axis (***Figure 1C and D***), suggesting that coordinated epithelial hypertrophy and villus expansion drive enlargement of the maternal intestine.

**Figure 1.**
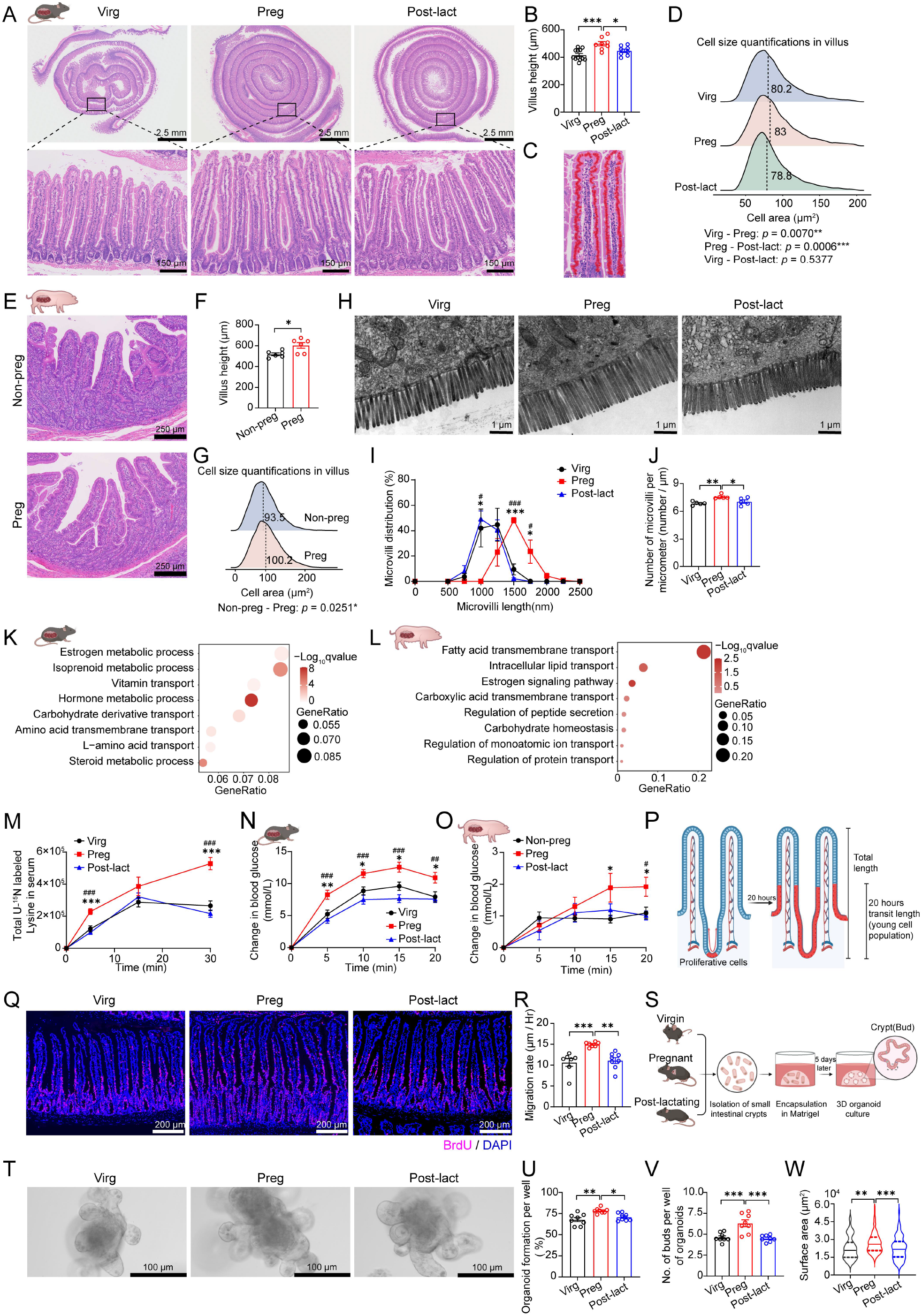
Pregnancy induces conserved intestinal remodeling across the mammalian species examined to enhance maternal nutrient absorption. (A) Representative hematoxylin and eosin (H&E)-stained sections of small intestine from virgin females (Virg), pregnant females at embryonic day 18.5 (Preg), and post-lactating females at day 35 (Post-lact). Scale bars, 2.5 mm (top) and 150 μm (bottom). (B) Quantification of the heights of small intestinal villi in Virg, Preg, and Post-lact females. (Virg, *n* = 12; Preg, *n* = 8; Post-lact, *n* = 8). (C) Representative H&E-stained section showing villus epithelial cell detection using QuPath. Scale bar, 50 μm. (D) Quantification of villus epithelial cell size in the jejunum among the three groups. (*n* = 8 per group). (E) Representative H&E-stained sections of the jejunum from non-pregnant (Non-preg) and late-pregnant (Preg) sows. Scale bar, 250 μm. (F and G) Quantification of jejunal villi height (F) and epithelial cell size (G) in Non-preg and Preg sows (n = 6 per group). (H) Representative electron microscopy (EM) images of jejunal enterocyte microvilli from Virg, Preg, and Post-lact females. Scale bar, 1 μm. (I and J) Distribution of microvilli lengths (I) and morphometric quantification of microvilli number per micrometer (J) based on EM images. (*n* = 5 per group). (K and L) Functional enrichment analysis of genes upregulated in the jejunum of pregnant mice (K) and pregnant sows (L) compared with non-pregnant controls. (M and N) Amino acid (M) (*n* = 5 per group) and glucose (N) (*n* = 11 per group) uptake rates following oral gavage in Virg, Preg, and Post-lact mice. (O) Glucose absorption rates following oral glucose administration in Non-preg, Preg, and Post-lact sows. (Non-preg, *n* = 11; Preg, *n* = 9; Post-lact, *n* = 7). (P) Schematic illustration of the BrdU tracing strategy. BrdU-labeled cells migrate along the villus axis over time following DNA synthesis labeling. (Q) Representative immunofluorescence analysis of BrdU-labeled intestinal epithelial cells after BrdU administration 20 hours before euthanasia. 30 randomly selected villi were measured per mouse, and epithelial migration rates were calculated. Scale bar, 200 μm. (R) Quantification of villus epithelial cell migration rates among Virg, Preg, and Post-lact females. (Virg, *n* = 7; Preg, *n* = 7; Post-lact, *n* = 8). (S) Schematic illustration of the intestinal organoid culture workflow. Small intestinal crypts isolated from Virg, Preg, and Post-lact females were cultured for 5 days. (T) Representative images of small intestinal organoids. Scale bar, 100 μm. (U and V) Quantification of organoid formation efficiency (U) and number of buds (V) after 5 days of culture (n = 4 mice per group; two wells analyzed per mouse). (W) Quantification of organoid surface area after 5 days of culture (n = 4 mice per group; 71, 113, and 87 organoids analyzed in the Virg, Preg, and Post-lact groups, respectively). Data are presented as means ± SEM. Statistical significance was determined by unpaired two-tailed Student’s *t*-test (F) or one-way ANOVA followed by Tukey’s multiple comparisons test. \**p* < 0.05, \*\**p* < 0.01, and \*\*\**p* < 0.001. For panels I, M, N, and O, ^#^*p* < 0.05, ^##^*p* < 0.01, and ^###^*p* < 0.001 (Preg vs. Post-lact).

To determine whether this adaptive response is conserved across mammals, we next analyzed jejunal tissues from non-pregnant and late-gestation sows. Similar to the changes observed in mice, pregnancy markedly increased villus height and epithelial cell size in the porcine jejunum compared with non-pregnant controls (***Figure 1E* *to* *G*; *Figure 1-supplementary figure 1D***). These conserved morphological adaptations across mice and pigs indicate that pregnancy-induced intestinal remodeling represents an evolutionarily conserved feature of mammalian gestation.

Microvilli form the apical brush border of small intestinal epithelial cells and increase the absorptive surface area by up to 25-fold (***Boyd & Parsons, 1969; Chevalier et al., 2015***). We therefore examined epithelial ultrastructure by transmission electron microscopy (TEM). Pregnancy significantly increased both microvillus length and density, thereby expanding the calculated absorptive surface area of individual enterocytes (***Figure 1H* *to* *J***). After weaning recovery, microvillus length largely returned to the level observed in virgin females (***Figure 1H* *to* *J***), further supporting the reversible nature of pregnancy-induced intestinal remodeling.

To identify the transcriptional programs accompanying these structural adaptations, we performed bulk RNA sequencing (RNA-seq) of jejunal tissues from virgin and late-gestation mice. Principal component analysis showed transcriptional separation between the two physiological states (***Figure 1-supplementary figure 1E***). Gene ontology (GO) analysis identified significant enrichment of biological processes related to amino acid transport, carbohydrate transport, vitamin absorption, transmembrane solute transport, and epithelial nutrient handling during pregnancy (***Figure 1K***). Consistent with these findings, transcriptomic analysis of jejunal tissues from pregnant sows showed concordant enrichment of nutrient transport and absorptive pathways (***Figure 1L*; *Figure 1-supplementary figure 1F***). Together, these results demonstrate that pregnancy induces conserved structural and transcriptional remodeling of the small intestine to increase nutrient absorption in both species examined.

We next asked whether these structural and transcriptional changes enhanced intestinal nutrient absorption function. Oral nutrient absorption assays showed significantly faster absorption of representative essential amino acids and monosaccharides in pregnant mice than in virgin controls (***Figure 1M and N***). After weaning recovery, nutrient absorption returned to levels comparable to those of virgin females (***Figure 1M and N***), consistent with the reversal of intestinal remodeling. We observed similar changes in pregnant sows, which exhibited accelerated nutrient absorption that also normalized after weaning recovery (***Figure 1O***). Together, these findings demonstrate that pregnancy induces coordinated structural, molecular, and functional remodeling of the maternal intestine to enhance nutrient absorption.

We next investigated whether pregnancy-induced expansion of the small intestinal epithelium is driven by increased ISC activity. Using 5-bromo-2′-deoxyuridine (BrdU) pulse-chase labeling (***Figure 1P***), we found that epithelial cell migration along the crypt–villus axis was significantly accelerated during pregnancy and largely returned to baseline after weaning recovery (***Figure 1Q and R***), indicating enhanced epithelial turnover. Consistent with these findings, intestinal crypts isolated from pregnant mice generated more organoids, displayed greater budding efficiency, and formed larger organoids than crypts from virgin or post-weaning mice (***Figure 1S* *to* *W***), demonstrating increased intrinsic regenerative capacity.

### Estradiol drives gestational intestinal remodeling independently of fetal demand

Gestational remodeling of the maternal small intestine is widely thought to arise from the increasing metabolic demands of fetal growth (***Casirola & Ferraris, 2003; Meyer & Caton, 2016; Overduin, Page, Young, & Gatford, 2025***). To determine whether this process is driven primarily by fetal-derived signals or maternal endocrine cues, we used a pseudopregnancy model, in which females undergo pregnancy-like hormonal changes in the absence of conceptuses. Pseudopregnant mice developed intestinal remodeling comparable to that of pregnant mice, including increased intestinal size, villus elongation, epithelial hypertrophy, and accelerated epithelial migration (***Figure 2A* *to* *E*; *Figure 2-supplementary figure 2A to C***). These findings show that intestinal remodeling proceeds in the absence of conceptuses, and that maternal physiological signals alone reproduce this adaptive response.

**Figure 2.**
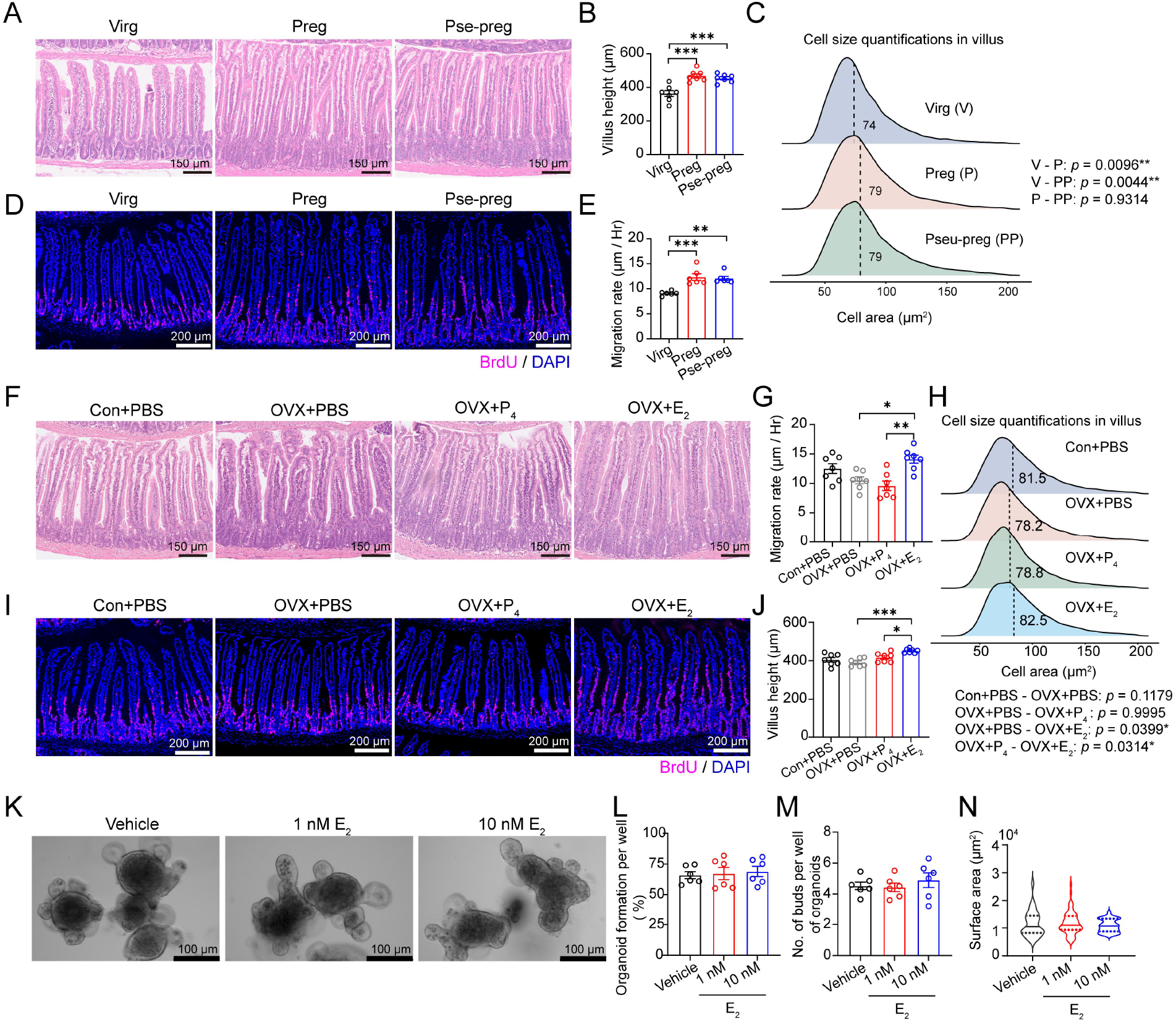
Estradiol drives gestational intestinal remodeling independently of fetal demand. (A–C) Representative images of small intestinal sections (A), quantification of villi heights (B), and villus epithelial cell size distributions (C) in virgin (Virg), pregnant (Preg), and pseudopregnant (Pse-preg) females. Scale bar, 150 μm. (*n* = 7 per group). (D and E) Representative immunofluorescence images of BrdU-labeled intestinal epithelial cells (D) and quantification of villus epithelial cell migration rates (E) in Virg, Preg, and Pse-preg females. Scale bar, 200 μm. (*n* = 6 per group). (F and G) Ovariectomized (OVX) mice were treated with estradiol (E2), progesterone (P4), or vehicle control for 10 days. Representative images of small intestinal villi (F) and quantification of villi heights (G). (*n* = 7 per group). (H) Quantification of villus epithelial cell size distributions using QuPath. (*n* = 7 per group). (I and J) Representative immunofluorescence images of BrdU-labeled intestinal epithelial cells (I) and quantification of villus epithelial cell migration rates (J). Scale bar, 200 μm. (*n* = 7 per group). (K) Representative images of small intestinal organoids following estradiol treatment. Scale bar, 100 μm. (L and M) Quantification of organoid formation efficiency (L) and number of buds (M). (n = 3 mice per group; two wells analyzed per mouse). (N) Quantification of organoid surface area (N) (n = 3 mice per group; 61, 61, and 55 organoids analyzed from the Vehicle, 1 nM, and 10 nM groups, respectively). Data are presented as means ± SEM. Statistical significance was determined by one-way ANOVA followed by Tukey’s multiple comparisons test. \**p* < 0.05, \*\**p* < 0.01, and \*\*\**p* < 0.001.

Pregnancy is accompanied by endocrine remodeling, characterized by progressive increases in circulating estradiol and progesterone, both of which have been implicated in maternal tissue adaptation. We therefore sought to identify the hormone responsible for gestational intestinal remodeling. Ovariectomized adult mice received physiological replacement of either estradiol or progesterone, and we subsequently examined intestinal morphology. Estradiol treatment markedly increased small intestinal length and weight and induced villus expansion that closely recapitulated the pregnancy phenotype (***Figure 2F* *to* *J*; *Figure 2-supplementary figure 2D to F***). In contrast, progesterone treatment produced minimal effects on intestinal morphology (***Figure 2F* *to* *J*; *Figure 2-supplementary figure 2D to F***). Together, these findings identify estradiol, and not progesterone, as an endocrine driver of pregnancy-induced intestinal remodeling.

To determine whether estradiol directly promotes the intrinsic regenerative capacity of the intestinal stem and progenitor cells, we cultured intestinal organoids in the presence of estradiol. Unexpectedly, estradiol did not alter organoid formation efficiency, budding capacity, or organoid size (***Figure 2K* *to* *N***), indicating that estradiol alone does not stimulate ISC activity in this system. These findings suggest that estradiol promotes pregnancy-induced intestinal remodeling indirectly by acting through niche-derived intermediary signals that couple systemic endocrine changes to ISC activation and epithelial renewal.

### Pregnancy-expanded Treg cells are required for intestinal remodeling

Given the established role of immune cells in epithelial regeneration and the immunological effects of estradiol (***Biton et al., 2018; Chakraborty et al., 2023; Forsyth, Jiwrajka, Lovell, Toothacre, & Anguera, 2024; Hoffmann, Liu, Seddu, & Klein, 2023; Lindemans et al., 2015***), we tested whether pregnancy-induced endocrine signals remodel the maternal intestine through specific immune cell populations. To identify these mediators, we performed single-cell RNA sequencing (scRNA-seq) of CD45^+^ lamina propria immune cells isolated from virgin and late-pregnant mice (***Figure 3A***). Analysis of 62,664 high-quality cells identified all major intestinal immune populations, including T cells, B cells, NK cells, innate lymphoid cells (ILCs), and myeloid cells (***Figure 3B; Figure 3-supplementary figure 3A and B***). Pregnancy extensively remodeled the intestinal immune landscape, with the most prominent changes occurring within the lymphoid compartment (***Figure 3C and D***). Milo analysis (***Dann, Henderson, Teichmann, Morgan, & Marioni, 2022***) revealed marked expansion of both T cells and ILCs, whereas Augur analysis (***Skinnider et al., 2021***) identified T cells as the immune population most responsive to pregnancy (AUC = 0.727) (***Figure 3C and D***). Consistent with these findings, transcriptional heterogeneity analysis also ranked T cells as the immune compartment undergoing the greatest gestational remodeling (***Figure 3E***). Together, these complementary analyses identified intestinal T cells as the immune population responding most strongly to pregnancy.

**Figure 3.**
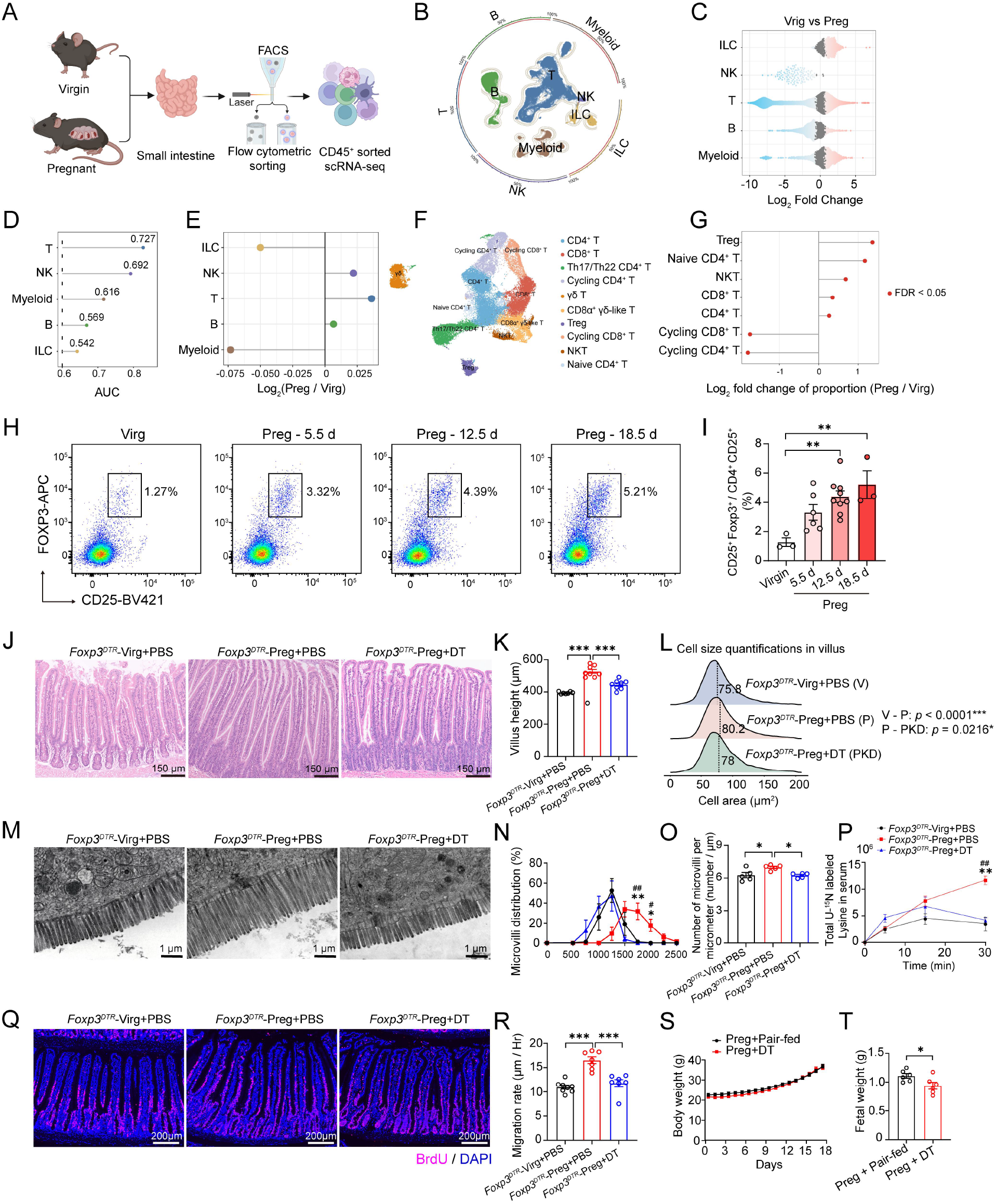
Pregnancy-expanded Treg cells are required for intestinal remodeling. (A) Schematic workflow of fluorescence-activated cell sorting (FACS) isolation of CD45^+^ immune cells from the small intestines of virgin (Virg) and pregnant (Preg) females. Three biological replicates were analyzed per group. CD45^+^ immune cells were pooled from two females per replicate. (B) UMAP visualization of 62,664 intestinal immune cells with major cell populations annotated based on marker gene expression. (C) Milo bee-swarm plot showing differential abundance of cellular neighborhoods across intestinal immune cell populations between Virg and Preg females. Significant neighborhoods with a 10% spatial false discovery rate (FDR) are highlighted. (D) Augur analysis showing the area under the curve (AUCs) scores of immune cell populations ranked by their sensitivity to pregnancy-associated transcriptional changes. (E) Transcriptional heterogeneity analysis showing log₂ (Preg/Virg) values across intestinal immune cell types. (F) UMAP visualization of intestinal T cells with major T cell subsets annotated according to characteristic marker gene expression profiles. (G) Log₂ fold change to the proportion of intestinal T cell subsets (Preg/Virg) identified via secondary clustering. Differential abundance was considered significant at FDR < 0.05. (H and I) Representative flow cytometry plots showing identification of small intestinal lamina propria Treg cells as CD45^+^CD4^+^CD25^+^Foxp3^+^ among live single CD45⁺ leukocytes (H), and quantification of the Treg cell proportion during pregnancy (I). (Virgin, *n* = 3; Preg - 5.5 d, *n* = 6; Preg - 12.5 d, *n* = 9; Preg - 18.5 d, *n* = 3). (J–L) Representative images of small intestinal sections (J), quantification of villi heights (K) (*n* = 8 per group), and villus epithelial cell size distribution (L) (*n* = 7 per group) in *Foxp3^DTR^-*Virg+PBS, *Foxp3^DTR^-*Preg+PBS, and *Foxp3^DTR^-*Preg+DT mice. Scale bar, 150 μm. (M–O) Representative electron microscopy images of jejunal enterocyte microvilli (M). Scale bar, 1 μm. Distribution of microvilli lengths (N) and morphometric quantification of microvilli numbers per micrometer (O). (*n* = 5 per group). (P) Amino acid uptake rates in *Foxp3^DTR^-*Virg+PBS, *Foxp3^DTR^-*Preg+PBS, and *Foxp3^DTR^-*Preg+DT mice. (*n* = 3 per group). (Q and R) Representative immunofluorescence images of BrdU-labeled intestinal epithelial cells (Q) and quantification of villus epithelial cell migration rates (R) in *Foxp3^DTR^*-Virg+PBS, *Foxp3^DTR^*-Preg+PBS, and *Foxp3^DTR^*-Preg+DT mice. Scale bar, 200 μm. (*Foxp3^DTR^-*Virg+PBS, *n* = 8; *Foxp3^DTR^-*Preg+PBS, *n* = 7; *Foxp3^DTR^-*Preg+DT, *n* = 7). (S and T) Pair-feeding experiment. Pair-fed mice received PBS and were restricted to the exact same food intake as the DT-treated group from embryonic day 11.5 (E11.5). Maternal body weight changes (S) and fetal weights (T). (*n* = 6 per group). Data are presented as means ± SEM. Statistical significance was determined by one-way ANOVA followed by Tukey’s multiple comparisons test. \**p* < 0.05, \*\**p* < 0.01, and \*\*\**p* < 0.001. For panels N and P, ^#^*p* < 0.05 and ^##^*p* < 0.01 (*Foxp3^DTR^-*Preg+PBS vs. *Foxp3^DTR^-*Preg+DT).

To define the T-cell populations responsible for this response, we performed secondary clustering of intestinal T cells. This analysis identified naive CD4^+^ T cells, conventional CD4^+^ and CD8^+^ T cells, Th17/Th22 cells, Treg cells, NKT cells, γδ T-cell subsets, and cycling CD4^+^ and CD8^+^ T cells based on their characteristic marker-expression profiles (***Figure 3F*; *Figure 3-supplementary figure 3C***). Among these subsets, Treg cells showed the most pronounced expansion during pregnancy (***Figure 3G***). Flow cytometric analysis independently confirmed a substantial increase in intestinal Treg-cell abundance during pregnancy (***Figure 3H and I***). These findings prompted us to investigate whether pregnancy-induced expansion of intestinal Treg cells is required for maternal intestinal remodeling.

To directly assess the functional requirement for Treg cells in pregnancy-associated intestinal remodeling, we transiently reduced Treg-cell abundance during late gestation using *Foxp3^DTR^* transgenic mice. To minimize the systemic and reproductive consequences associated with profound Treg-cell ablation, we adopted a low-dose diphtheria toxin (DT) regimen that induced partial, rather than complete, depletion of Treg cells. Under this regimen, pregnant mice showed no overt maternal illness or substantial pregnancy loss (***Figure 3-supplementary figure 4A and B***), indicating that the intervention was well tolerated during late gestation. Partial reduction of Treg cells nevertheless markedly impaired pregnancy-induced intestinal remodeling (***Figure 3J* *to* *O*; *Figure 3-supplementary figure 4C to E***). Compared with pregnant controls, DT-treated mice exhibited reduced villus height, smaller epithelial cells, and diminished microvillus elongation, resulting in marked attenuation of pregnancy-associated intestinal expansion (***Figure 3J* *to* *O***). Consistent with these structural abnormalities, pregnancy-induced enhancement of nutrient absorption was largely abolished following Treg-cell reduction (***Figure 3P***). BrdU pulse-chase analysis further demonstrated significantly reduced epithelial migration along the crypt–villus axis, indicating impaired intestinal stem-cell–driven epithelial renewal (***Figure 3Q and R***). To assess the contribution of reduced maternal food intake, we performed pair-feeding experiments. Despite comparable changes in maternal body weight, pair-fed controls showed significantly better intestinal phenotypes than DT-treated mice and did not exhibit the reduction in fetal weight observed in DT-treated mice (***Figure 3S* *to* *T*; *Figure 3-supplementary figure 4F to I***). Together with the nutrient absorption data, these findings suggest that Treg cells contribute to maternal intestinal remodeling and fetal growth beyond the effects of reduced food intake. Together, these findings support a functional requirement for Treg cells in orchestrating maternal intestinal remodeling during pregnancy.

### Estradiol signaling in Treg cells enables intestinal remodeling during pregnancy

Having identified intestinal Treg cells as the immune population most responsive to pregnancy, we next investigated whether estradiol directly regulates Treg-cell function through estrogen receptor signaling. *Esr1* expression was highly enriched in immune cells (***Chakraborty et al., 2023; Cunningham & Gilkeson, 2011; Kim et al., 2019; Kovats, 2015***), suggesting that estradiol may directly regulate Treg-cell function through estrogen receptor α (ERα). To determine the physiological importance of ERα signaling in Treg cells, we generated mice with a Treg-specific deletion of *Esr1* (*Esr1^Foxp3^*) and examined intestinal adaptation during pregnancy. Compared with pregnant *Esr1^fl/fl^* littermates, pregnant *Esr1^Foxp3^* mice exhibited a marked impairment in pregnancy-induced intestinal remodeling. Treg-specific deletion of ERα significantly reduced villus elongation, epithelial hypertrophy, and microvillus extension, resulting in a substantial loss of intestinal absorptive surface area (***Figure 4A* *to* *F*; *Figure 4-supplementary figure 5A to C***). Consistent with these structural defects, the pregnancy-associated increase in intestinal nutrient absorption was largely abolished in *Esr1^Foxp3^* mice (***Figure 4G***), demonstrating that ERα-mediated estrogen signaling in Treg cells is essential for establishing the enhanced absorptive capacity of the maternal intestine during pregnancy.

**Figure 4.**
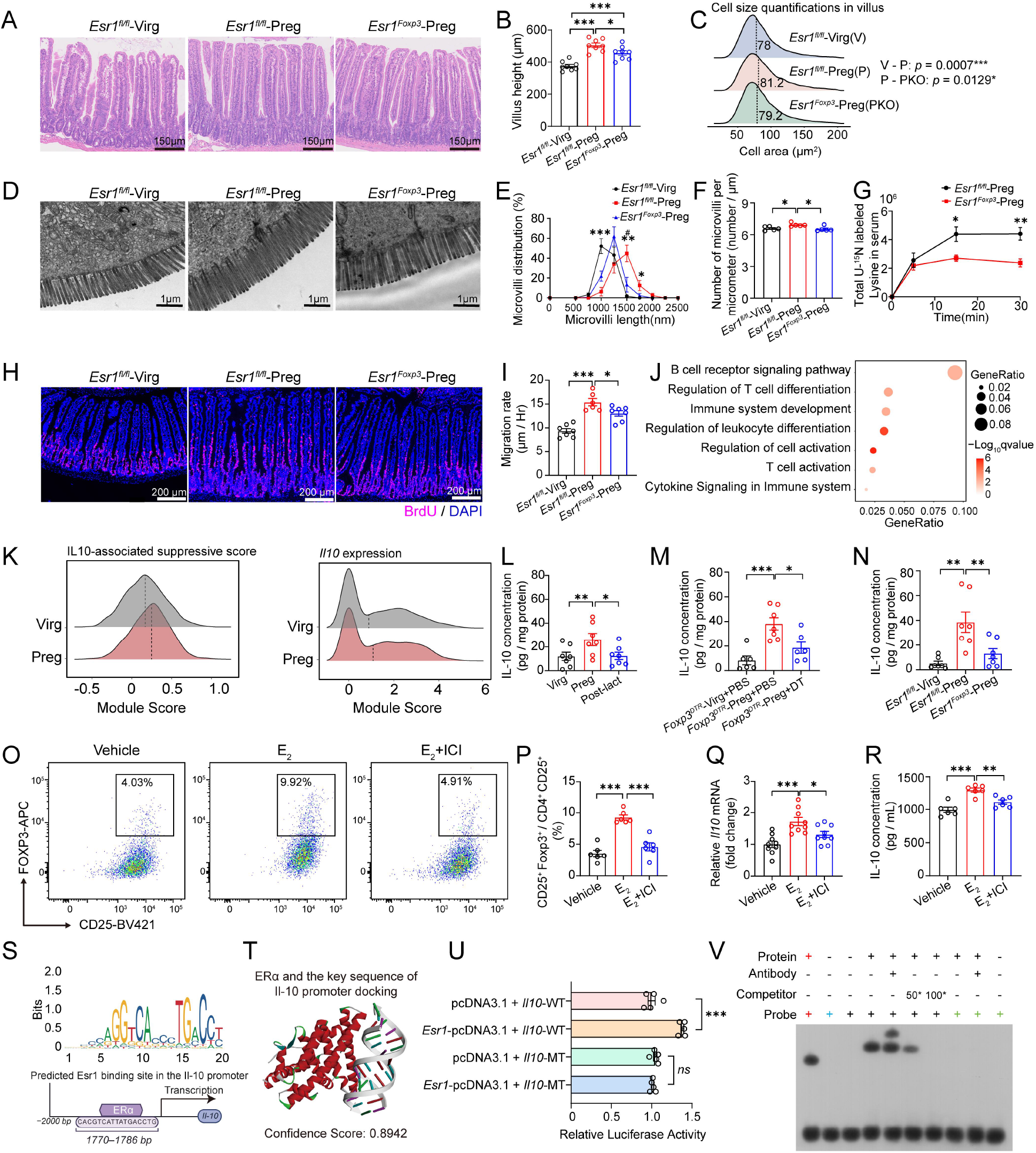
Estradiol signaling in Treg cells enables intestinal remodeling during pregnacy. (A–C) Representative images of small intestinal sections (A), quantification of villi heights (B) (*n* = 8 per group), and villus epithelial cell size distributions (C) (*n* = 7 per group) in *Esr1^fl/fl^*-Virg, *Esr1^fl/fl^*-Preg, and *Esr1^Foxp3^*-Preg mice. Scale bar, 150 μm. (D–F) Representative electron microscopy images of jejunal enterocyte microvilli (D). Scale bar, 1 μm. Distribution of microvilli lengths (E) and morphometric quantification of microvilli numbers per micrometer (F). (*n* = 5 per group). (G) Amino acid uptake rates in *Esr1^fl/fl^*-Virg, *Esr1^fl/fl^*-Preg, and *Esr1^Foxp3^*-Preg mice. (*n* = 4 per group). (H and I) Representative immunofluorescence images of BrdU-labeled intestinal epithelial cells (H) and quantification of villus epithelial cell migration rates (I). Scale bar, 200 μm. (*Esr1^fl/fl^*-Virg, *n* = 7; *Esr1^fl/fl^*-Preg, *n* = 6; *Esr1^Foxp3^*-Preg, *n* = 7). (J) Functional enrichment analysis of genes upregulated in Treg cells from pregnant mice compared with virgin mice. (K) Module score analysis of IL-10–associated suppressive signatures and *Il10* expression in Treg cells from pregnant and virgin mice. (L–N) Quantification of intestinal IL-10 levels in Virg, Preg, and Post-lact females (L) (*n* = 7 per group); *Foxp3^DTR^-*Virg+PBS, *Foxp3^DTR^-*Preg+PBS, and *Foxp3^DTR^-*Preg+DT mice (M) (*Foxp3^DTR^-*Virg+PBS, *n* = 6; *Foxp3^DTR^-*Preg+PBS, *n* = 7; *Foxp3^DTR^-*Preg+DT, *n* = 6); and *Esr1^fl/fl^*-Virg, *Esr1^fl/fl^*-Preg, and *Esr1^Foxp3^*-Preg mice (N) (*n* = 7 per group) as measured by ELISA. (O and P) Primary CD4^+^ T cells isolated from virgin mice were treated with estradiol or the estrogen receptor antagonist ICI 182,780 (ICI). Representative flow cytometry plots showing Treg cell identification as CD45^+^CD4^+^CD25^+^Foxp3^+^ cells (O) and quantification of Treg cell frequency (P). (*n* = 6 per group). (Q and R) Relative mRNA expression of *Il10* (Q) (*n* = 9 per group) and IL-10 concentrations in culture supernatants following treatment (R) (*n* = 6 per group). (S) Transcription factor motif prediction of ERα-binding sites within the *Il10* promoter region. (T) Predicted DNA–protein docking interaction between ERα and the *Il10* promoter. (U) Luciferase reporter assay assessing ERα-dependent transcriptional activation of the *Il10* promoter. (V) EMSA validation of ERα-DNA binding. Lanes: 1, Positive control; 2, Negative control; 3, WT probe alone; 4, WT + ERα; 5, WT + ERα + Anti-Halo Ab (supershift); 6, WT + ERα + 50* unlabeled WT competitor; 7, WT + ERα + 100* unlabeled WT competitor; 8, MT + ERα; 9, MT + ERα + Anti-Halo Ab; 10, MT probe alone. (Ab, antibody; WT, wild-type; MT, mutant). Data are presented as means ± SEM. Statistical significance was determined by an unpaired two-tailed Student’s t test (G) or one-way ANOVA followed by Tukey’s multiple comparisons test. \**p* < 0.05, \*\**p* < 0.01, and \*\*\**p* < 0.001.

BrdU pulse-chase analysis further showed that epithelial cell migration along the crypt–villus axis was significantly slower in pregnant *Esr1^Foxp3^* mice than in pregnant littermate controls (***Figure 4H and I***), indicating ISC-driven renewal is impaired during gestation. These defects closely phenocopied those observed following transient Treg-cell reduction, which is consistent with estradiol acting on intestinal remodeling through ERα signaling in Treg cells, although the phenocopy does not exclude contributions from other estrogen-responsive populations.

We next sought to define the downstream molecular program activated by ERα signaling in Treg cells during pregnancy. Differential gene expression analysis showed extensive pregnancy-induced transcriptional remodeling, with significant enrichment of immune regulatory pathways (***Figure 4***J). Among the differentially regulated genes, pregnancy markedly increased both the proportion of *Il10*-expressing Treg cells and the fraction of Treg cells with high IL-10 pathway activity (***Figure 4K***). These complementary analyses identified IL-10 as a downstream effector of estradiol-responsive Treg cells during pregnancy. To test this relationship in vivo, we measured intestinal IL-10 expression following disruption of the pathway by either transiently reducing Treg cells or Treg-specific deletion of *Esr1*. Pregnancy markedly increased IL-10 expression in the small intestine, whereas both interventions almost completely abolished this induction (***Figure 4L* *to* *N***). Together, these findings show that estradiol–ERα signaling in intestinal Treg cells is required for the pregnancy-associated rise in intestinal IL-10, measured here in whole tissue.

To test this relationship directly, we differentiated naive CD4^+^ T cells into Treg cells in vitro and treated them with estradiol in the presence or absence of the estrogen receptor antagonist ICI 182,780. Estradiol significantly enhanced Treg-cell differentiation and markedly increased *Il10* expression and IL-10 secretion, whereas pharmacological inhibition of estrogen receptor signaling completely abolished these effects (***Figure 4O* *to* *R***). These results demonstrate that estradiol promotes Treg-cell differentiation and IL-10 production through estrogen receptor signaling, contributing to the pregnancy-associated increase in intestinal IL-10.

In silico analysis using the hIFtarget databases identified a predicted ERα-binding motif within the *Il10* promoter (***Figure 4S***). DNA–protein docking simulations predicted binding of ERα to this region (***Figure 4T***). To test this prediction, we performed *Il10* promoter luciferase reporter assays. ERα overexpression significantly increased *Il10* promoter activity, whereas mutation of the predicted ERα-binding motif partially attenuated this effect (***Figure 4U***). To further validate ERα binding to the *Il10* promoter, we performed electrophoretic mobility shift assays (EMSA). A specific super-shift band was observed only in the presence of ERα protein, the WT probe, and an ERα antibody (***Figure 4V***). This band disappeared when either a mutant (MT) probe was used or a competitive unlabeled probe was added, confirming specific ERα–DNA interaction (***Figure 4V***). These results demonstrate that ERα can directly binds to the *Il10* promoter and promotes its transcription.

### Treg-dependent IL-10 directly stimulates intestinal stem-cell activity to drive gestational intestinal remodeling

Having identified IL-10 as a downstream effector of estradiol-responsive Treg cells, we next examined whether IL-10 directly regulates ISC-dependent epithelial regeneration. Treatment of intestinal organoids with recombinant IL-10 significantly increased organoid-forming efficiency in a dose-dependent manner (***Figure 5A and B***). IL-10 also markedly enhanced organoid budding, consistent with increased ISC-driven epithelial growth (***Figure 5C***). Consistent with these findings, 5-ethynyl-2′-deoxyuridine (EdU) incorporation showed an increase in proliferating epithelial cells following IL-10 treatment (***Figure 5D* *to* *H***). Pharmacological blockade of the IL-10 receptor α (IL-10Rα/CD210) completely abolished these effects (***Figure 5D* *to* *H***), demonstrating that IL-10 directly promotes intestinal epithelial regeneration through IL-10R signaling.

**Figure 5.**
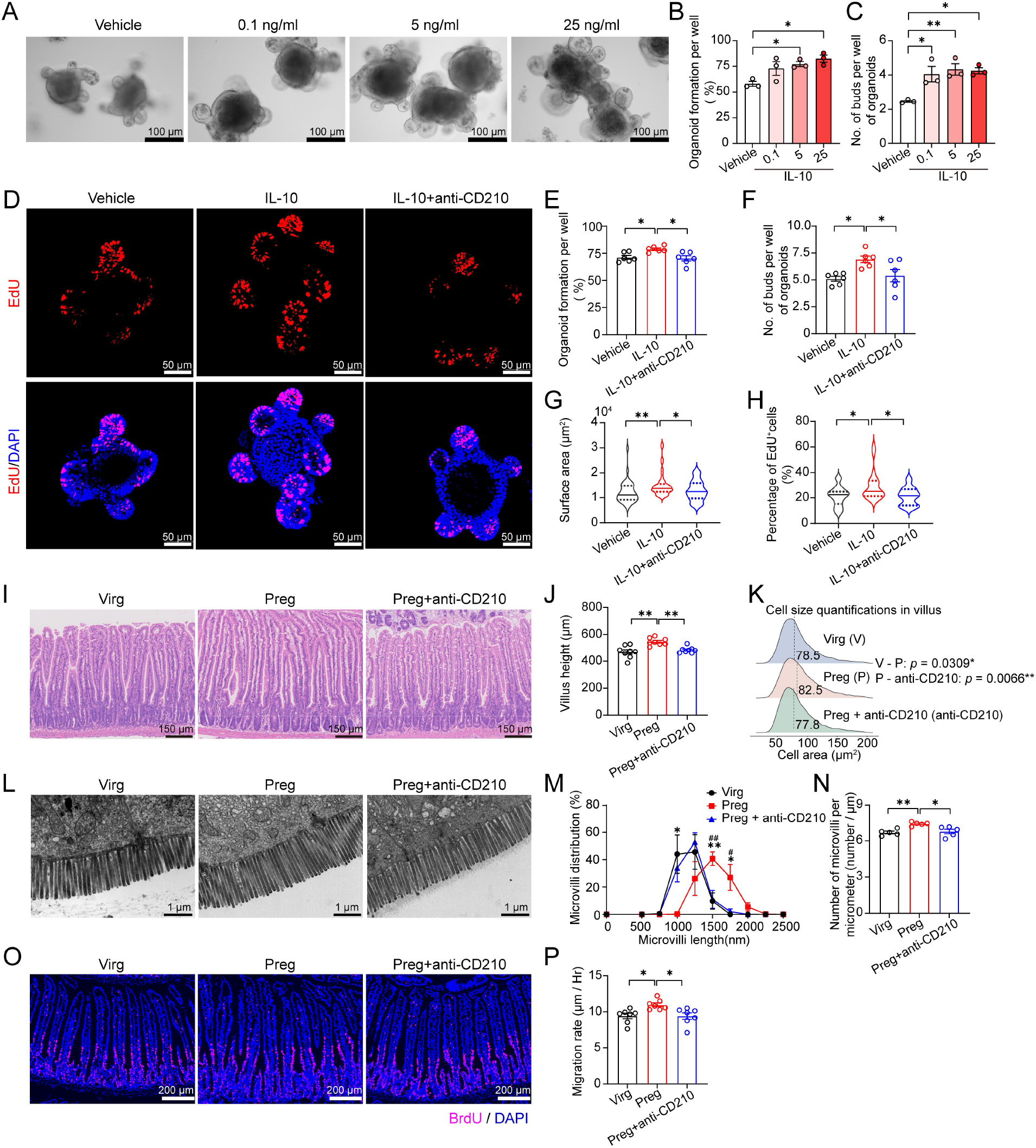
Treg-dependent IL-10 directly stimulates intestinal stem-cell activity to drive gestational intestinal remodeling. (A–C) Representative images of small intestinal organoids following IL-10 treatment (A), quantification of organoid formation efficiency (B), and number of buds (C). Scale bar, 100 μm. (*n* = 3 mice per group). (D and H) Representative images of EdU staining in small intestinal organoids following treatment with IL-10 and a blocking antibody against the IL-10 receptor (anti-CD210) (D), and quantification of EdU^+^ cell percentage (H). Scale bar, 50 μm. (n = 3 mice per group; 20 organoids analyzed from each group). (E and F) Quantification of organoid formation efficiency (E) and number of buds (F) following IL-10R inhibition. (G) Quantification of organoid surface area following IL-10R inhibition. (n = 3 mice per group; 40, 45, and 45 organoids analyzed from the Vehicle, IL-10, and IL-10+anti-CD210 groups, respectively). (I–K) Representative images of small intestinal sections from virgin (Virg), pregnant (Preg), and pregnant mice treated with anti-CD210 (Preg+anti-CD210) (I). Quantification of villI heights (J) (*n* = 8 mice per group), and villi epithelial cell size distribution (K) (*n* = 7 mice per group). Scale bar, 150 μm. (L–N) Representative electron microscopy images of jejunal enterocyte microvilli (L). Scale bar, 1 μm. Distribution of microvilli lengths (M) and morphometric quantification of microvilli numbers per micrometer (N). (*n* = 5 mice per group). (O and P) Representative immunofluorescence images of BrdU-labeled intestinal epithelial cells (O) and quantification of villus epithelial cell migration rates (P). Scale bar, 200 μm. (*n* = 7 mice per group). Data are presented as means ± SEM. Statistical significance was determined by one-way ANOVA followed by Tukey’s multiple comparisons test. \**p* < 0.05, \*\**p* < 0.01, and \*\*\**P* < 0.001. For panel M, ^#^*p* < 0.05 and ^##^*p* < 0.01 (Preg vs. Preg+anti-CD210).

We next determined whether IL-10 signaling is required for pregnancy-induced intestinal remodeling in vivo. Pregnant mice received an IL-10R-blocking antibody at E12.5, coinciding with the rapid phase of gestational intestinal expansion. IL-10R blockade markedly impaired intestinal remodeling without altering the progression of pregnancy. Histological analysis showed that inhibition of IL-10 signaling significantly reduced villus elongation, epithelial hypertrophy, and microvillus extension compared with pregnant control mice (***Figure 5I* *to* *N*; *Figure 5-supplementary figure 6A to C***), resulting in a marked reduction in pregnancy-associated absorptive surface area. BrdU pulse-chase analysis further demonstrated that IL-10R blockade significantly slowed epithelial cell migration along the crypt–villus axis (***Figure 5O and P***), consistent with impaired ISC-driven epithelial renewal during pregnancy. Collectively, these findings identify IL-10 as a molecular mediator linking estradiol-dependent activation of Treg cells to ISC function.

### The estradiol–Treg–IL-10 axis couples maternal intestinal adaptation with fetal growth and is conserved in humans

Having established that estradiol-responsive Treg cells promote intestinal remodeling through IL-10-dependent activation of ISCs, we next investigated whether disruption of this pathway impairs maternal adaptation and fetal development during pregnancy. To address this question, we evaluated pregnancy outcomes in mice with Treg-specific deletion of ERα. Compared with pregnant *Esr1^fl/fl^* littermates, pregnant *Esr1^Foxp3^* females exhibited significantly reduced gestational weight gain throughout pregnancy (***Figure 6A***).They also accumulated significantly less maternal adipose tissue during late gestation (***Figure 6B***), indicating impaired maternal metabolic adaptation. These maternal defects were accompanied by impaired fetal growth. Fetuses from *Esr1^Foxp3^* dams were significantly smaller and had lower birth weights than those from littermate controls (***Figure 6C and D***). Given the marked impairment in maternal intestinal remodeling observed in *Esr1^Foxp3^* mice, these findings suggest that disruption of the estradiol–Treg axis may contribute to limited maternal nutrient acquisition, potentially impacting nutrient availability to support fetal growth.

**Figure 6.**
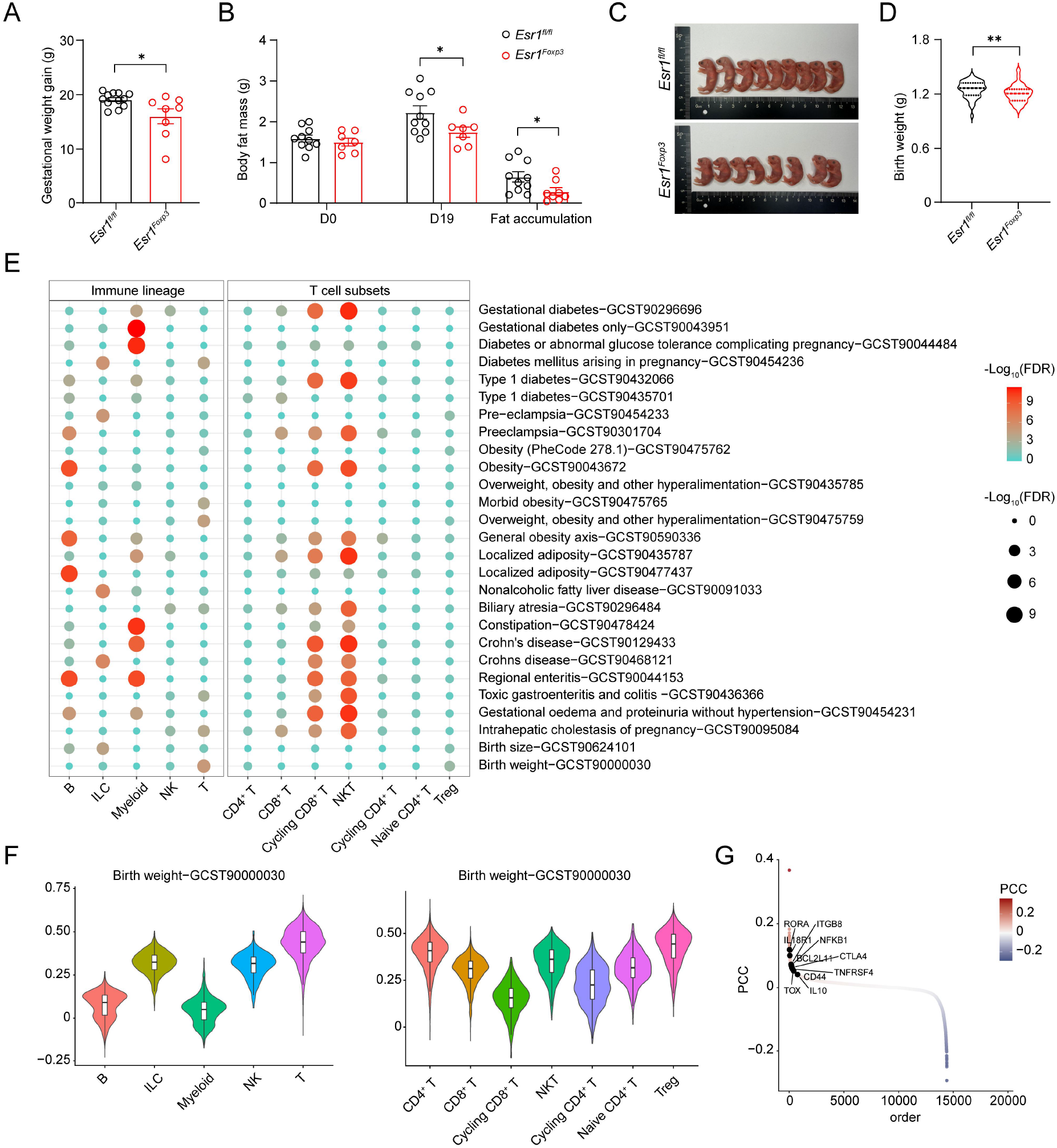
The estradiol–Treg–IL-10 axis couples maternal intestinal adaptation with fetal growth and is conserved in humans. (A–D) Quantification of gestational weight gain (A) (*Esr1^fl/fl^*, *n* = 12; *Esr1^Foxp3^*, *n* = 8), maternal fat mass from gestational days 0 to 19 and gestational fat accumulation (B) (*Esr1^fl/fl^*, *n* = 10; *Esr1^Foxp3^*, *n* = 7), representative images of fetuses (C), and birth weight of pups (D) (*Esr1^fl/fl^* and *Esr1^Foxp3^* dams total pups analyzed: *n* = 91 and 65, respectively) from *Esr1^fl/fl^* and *Esr1^Foxp3^* mice. (E) Bubble plot showing the associations between five immune lineages and five T subtypes with the 27 human complex traits related to pregnancy. (F) Trait-relevant score (TRS) depicting the correlations between five immune lineages and seven T subtypes with human birth weights. (G) Birth weight-relevant genes (human complex trait) were ranked by the Pearson correlation coefficient (PCC) using scPagwas across all individual Treg cells. Data are presented as means ± SEM. Statistical significance was determined by unpaired two-tailed Student’s t test. * *p* < 0.05, ** *p* < 0.01.

To assess the potential relevance of this regulatory pathway to human pregnancy, we next integrated our intestinal single-cell transcriptomic dataset with 27 human pregnancy-related genome-wide association studies (GWAS) using the scPagwas framework (***Figure 6-supplementary table 1***). Trait–cell type enrichment analysis showed that, among the intestinal immune populations profiled, pregnancy-associated traits scored highest in NK-cell and T-cell lineages (***Figure 6E***). Among the five intestinal T-cell subsets, Treg cells consistently showed the highest trait-relevance scores for birth weight and multiple pregnancy-associated metabolic traits (***Figure 6F***), suggesting that Treg-cell transcriptional programs broadly contribute to maternal metabolic adaptation and fetal development.

To further define the molecular basis of these associations, we prioritized genes contributing to birth-weight-associated transcriptional signatures in intestinal Treg cells. Network-based trait prioritization identified *Il10* as the top candidate, together with multiple downstream regulatory genes, including *Ctla4*, *Rora*, *Tox*, *Tnfrsf4*, *Nfkb1, Il18r1, Bcl2l11, Itgb8*, and *Cd44*, as major contributors to birth-weight-associated transcriptional programs (***Figure 6G***). These genes were upregulated in small intestinal Treg cells during pregnancy (***Figure 6G***), which is consistent with a conserved IL-10-centered gene set associated with fetal growth across the two species. Collectively, the concordance between the human genetic associations and the mouse experiments is consistent with physiological relevance and conservation of this endocrine–immune pathway during mammalian pregnancy.

## Discussion

Pregnancy requires coordinated physiological adaptations that enable the mother to meet the rapidly increasing nutritional demands of the developing fetus while maintaining maternal homeostasis. Although previous studies have largely focused on metabolic adaptations in the liver, adipose tissue, pancreas, and placenta (***Falick Michaeli et al., 2015; Mor et al., 2017; Sanghavi & Rutherford, 2014; Trowsdale & Betz, 2006; Yang et al., 2025; Yu et al., 2024***), the maternal intestine, the site of nutrient absorption, has received comparatively little attention. Here, we identify the maternal small intestine as an under-characterized target of gestational adaptation and show that pregnancy induces a conserved physiological growth and expansion program in mice and pigs. This program is characterized by coordinated villus expansion, accelerated epithelial renewal, transcriptional activation of nutrient transport pathways, and enhanced absorptive capacity. Our cross-species analyses further suggest that intestinal remodeling Our cross-species analyses further suggest that intestinal remodeling is an evolutionarily conserved adaptation that optimizes maternal nutrient acquisition to support fetal growth.

Pregnancy has traditionally been viewed through the lens of immune adaptations that establish and maintain maternal–fetal tolerance (***Aghaeepour et al., 2017; Erlebacher, 2013; Mor et al., 2017***). Our findings expand this paradigm by demonstrating that pregnancy-induced immune remodeling also acts on maternal organ function. Although Treg-cell expansion is widely recognized for its essential role in maintaining maternal–fetal immune tolerance (***Aluvihare et al., 2004; Li et al., 2025; Rowe et al., 2012; Samstein et al., 2012***), we show that gestational estradiol directly activates ERα signaling in intestinal Treg cells to induce a remodeling-associated transcriptional program centered on IL-10 production. Aligning with the emerging view that immunity acts as an active regulator of maternal physiology, (***Corral & Belkaid, 2026***) our study demonstrates that pregnancy repurposes immune tolerance pathways, extending the role of Tregs beyond immune suppression, to actively coordinate maternal tissue remodeling. More broadly, our findings provide a mechanistic framework through which systemic endocrine signals reprogram immune-cell function to alter adult stem-cell activity during physiological tissue remodeling.

Our findings also expand the biological functions of IL-10 beyond its well-established role as an anti-inflammatory cytokine. Although IL-10 is essential for maintaining intestinal immune homeostasis and protecting against inflammatory bowel disease (***Kühn, Löhler, Rennick, Rajewsky, & Müller, 1993; Moore, de Waal Malefyt, Coffman, & O’Garra, 2001; Rubtsov et al., 2008; Shouval et al., 2014***), its direct action on epithelial regeneration under physiological conditions has remained undefined. Using complementary organoid, genetic, and *in vivo* IL-10 receptor-blockade approaches, we demonstrate that Treg-dependent IL-10 directly promotes ISC activity and epithelial renewal during pregnancy. Together with recent studies showing that immune-derived cytokines act on tissue-specific stem cells in the skin, lung, and intestine (***Biton et al., 2018; Cohen et al., 2025; Glisinski et al., 2020; Gronke et al., 2019; Lindemans et al., 2015; Z. Liu et al., 2022; Mathur et al., 2019; Sullivan et al., 2021; Tadokoro et al., 2014***), our findings support a broader model in which immune cells actively instruct regenerative programs rather than simply responding to tissue injury. Importantly, we propose that this Treg–IL-10 axis does not act in isolation but represents a complementary layer within a broader endocrine– immune–epithelial remodeling network. Recent elegant studies have demonstrated that RANK-RANKL signaling provides an epithelial-intrinsic execution program for villus expansion (***Onji et al., 2025***), while eosinophils direct goblet cell differentiation (***Huang et al., 2026***). We propose that these pathways are not mutually exclusive; rather, the RANK-RANKL axis dictates the intrinsic capacity for epithelial expansion, whereas the estradiol–Treg–IL-10 axis establishes an immunotolerant microenvironment that permits and coordinates this extensive tissue remodeling safely and efficiently. Thus, pregnancy represents a physiological setting in which immune-mediated stem-cell regulation is repurposed to meet increased metabolic demands instead of repairing damaged tissue.

Our study further demonstrates that maternal intestinal adaptation is functionally coupled to fetal development. Treg-specific disruption of estradiol signaling impaired maternal intestinal remodeling, reduced nutrient absorption, compromised maternal metabolic adaptation, and ultimately restricted fetal growth. Consistent with these findings, integration of human genome-wide association studies implicated Treg-cell transcriptional programs, particularly the IL-10 regulatory network, as being associated with birth weight and multiple pregnancy-related metabolic traits. Although these human genetic associations do not establish causality, their concordance with mechanistic studies in mice, conserved intestinal remodeling in pigs, and physiological observations in pregnant women supports the translational relevance of this pathway. Together, these findings suggest that impaired maternal intestinal adaptation may represent an underrecognized mechanism contributing to fetal growth restriction and other pregnancy complications associated with inadequate maternal nutrient supply. Overall, our findings identify an estradiol–ERα–Treg–IL-10 axis that links maternal immune adaptation to intestinal remodeling during pregnancy. The conserved morphological adaptations observed in mice and pigs, together with human pregnancy GWAS analyses, support the potential cross-species relevance of this pathway and provide a mechanistic framework for how pregnancy coordinates maternal nutrient acquisition with fetal growth.

### Limitations of the study

Our study has several limitations. Although IL-10 is a downstream effector of estradiol-responsive Treg cells, additional pathways may also contribute to pregnancy-induced intestinal remodeling, and the signaling route by which IL-10 acts on intestinal stem cells remains to be elucidated. Furthermore, while our data indicate that Tregs are a major source of IL-10 during pregnancy, we cannot completely exclude potential contributions from other immune cell populations (such as macrophages) within the intestinal microenvironment. Moreover, acute Treg reduction using the *Foxp3^DTR^* model, even under a low-dose regimen that induced only partial depletion, cannot completely exclude secondary systemic effects associated with transient disruption of immune homeostasis during pregnancy. Importantly, the consistent phenotype observed in mice with Treg-specific deletion of *Esr1* independently supports a cell-autonomous requirement for estradiol signaling in Treg cells. Prospective human studies will also be required to establish the clinical relevance of this pathway.

## Materials and methods

### Mice

Female C57BL/6J mice (8 weeks old) were obtained from Vital River Laboratory Animal Technology (Beijing, China). *Foxp3^DTR^* mice (strain no. 016958) and *Esr1^fl/fl^* (strain no. 032173) mice were obtained from The Jackson Laboratory. Treg-specific *Esr1-*deficient mice (*Esr1^Foxp3^*) were generated by crossing *Esr1^fl/fl^* mice with *Foxp3-Cre* mice (Cyagen, C001467). Mice were maintained under specific pathogen-free conditions at 20–22°C on a 12-hour light/dark cycle and had *ad libitum* access to food and water.

For timed pregnancy experiments, female mice were housed with proven fertile males at a 1:1 ratio. The presence of a vaginal plug was designated as embryonic day 0.5 (E0.5). Post-lactation day 0 was defined as the day of pup removal at lactation day 21.

Pseudopregnancy was induced by mating females with vasectomized males, and plug-positive females were designated as pseudopregnant day 0.5. For the pseudopregnancy study, pregnant and pseudopregnant mice were analyzed at day 15.5. For the hormone supplementation study, bilateral ovariectomy was performed on adult females, while sham-operated mice underwent identical procedures without ovary removal. After a 2-week recovery period, the mice received daily intraperitoneal injections of 17β-estradiol (E_2;_ Selleck, S1709) at 1.5 μg, progesterone (P_4;_ Selleck, S1705) at 500 μg, or vehicle control for 10 consecutive days. For transient Treg-cell reduction, *Foxp3^DTR^* mice received intraperitoneal injections of diphtheria toxin (10 ng/g body weight) every two days from E11.5 to E17.5. To block IL-10R signaling, pregnant mice were intraperitoneally injected with anti-mouse IL-10Rα/CD210 antibody (MCE, HY-P990228) or vehicle control at E12.5 at a dose of 1 mg per mouse. Intestinal tissues were collected at E18.5 for subsequent analyses.

Mice were fasted for 5 hours before tissue collection. The small intestine was excised, weighed, and measured for length. Duodenal and jejunal tissues were prepared as Swiss rolls for histological analysis. Jejunal samples for transmission electron microscopy were immediately fixed in electron microscopy fixative, and additional tissues were snap-frozen in liquid nitrogen and stored at −80°C.

All animal procedures involving mice were approved by the Animal Ethics and Welfare Committee of Sichuan Agricultural University (Approval No. 20220138).

### Pigs

Twelve-month-old female pigs were used for validation studies. Jejunal tissues were collected from nonpregnant pigs and late-gestation pigs at gestational day 105. After euthanasia, jejunum segments were immediately collected, rinsed with sterile PBS, and divided for subsequent analyses. Samples for histological analysis were fixed in 4% paraformaldehyde, whereas samples for RNA sequencing were snap-frozen in liquid nitrogen and stored at −80°C until further processing. All procedures involving pigs were approved by the same committee (Approval No. 20230485).

For all animal experiments in this study (involving both mice and pigs), animals were randomly assigned to experimental groups prior to any treatment.

### Cell culture

For primary T cell culture, naive CD4^+^ T cells were isolated from mouse spleens using a magnetic bead-based isolation kit (Miltenyi Biotec, 130-104-453). Cells were activated on anti-CD3ε-coated plates (BD Biosciences, 553057) in RPMI-1640 supplemented with anti-CD28 antibody (BD Biosciences, 553294), recombinant IL-2 (R&D Systems, 402-ML-020), and TGF-β (R&D Systems, 240-B-002). To evaluate estradiol signaling, cells were treated with vehicle, estradiol (E2), or the estrogen receptor antagonist ICI-182780 (Selleck, S1191) before E2 stimulation.

For intestinal organoid culture, proximal small intestines were collected, flushed with cold DPBS containing 2% penicillin–streptomycin, opened longitudinally, and cut into 6–7 mm segments. Tissues were washed thoroughly and incubated in cold DPBS containing 5 mM EDTA for 30 min to release intestinal crypts. Crypt-containing fractions were collected by serial filtration and centrifugation. Isolated crypts were resuspended in Matrigel (bioGenous, M315077) and embedded as 50 μL Matrigel domes in 24-well plates. After Matrigel polymerization, complete organoid culture medium (bioGenous, K2001-MI) was added. Organoids were cultured at 37°C in a humidified incubator with 5% CO₂, and the medium was replaced every 48 h. Primary organoids isolated from virgin, pregnant, and post-lactating mice were used for phenotypic analyses. All subsequent experiments were performed using organoids after at least two passages.

### Organoid hormone and cytokine stimulation assays

To evaluate the direct effects of estradiol on intestinal organoids, organoids were treated with E_2 a_t indicated concentrations (vehicle, 1 nM, and 10 nM) for 24 hours.

To assess IL-10-mediated effects, organoids were treated with recombinant mouse IL-10 protein (Abcam, ab281800) at 0.1, 5, and 25 ng/ml for 24 hours. For IL-10 receptor blockade, organoids were pretreated with anti-mouse IL-10Rα/CD210 antibody for 1 hour before IL-10 stimulation (25 ng/ml) and subsequently cultured for an additional 24 hours. Vehicle-treated organoids served as controls. Organoids were collected for downstream analyses.

### Oral nutrient absorption assays

For glucose absorption assays, mice and pigs were fasted for 5 hours and 12 hours, respectively, before receiving oral glucose gavage. Glucose was administered at 1.25 mg/g body weight (mice) and 1 g/kg body weight (pigs). Blood samples were collected from the tail vein of mice and the ear vein of pigs at the indicated time points after gavage. Blood glucose levels were measured using a glucometer (Sinocare Gold-Accu).

For amino acid absorption assays, mice were fasted for 5 hours before oral administration of stable isotope-labeled essential amino acids by gavage as previously described (***Lee et al., 2017***). Tail vein blood was collected at the indicated time after gavage. Serum metabolites were extracted by methanol precipitation and analyzed by liquid chromatography–mass spectrometry (LC–MS) using an Agilent 1290 II UPLC coupled to a SCIEX 5600+ quadrupole time-of-flight mass spectrometer. Stable isotope enrichment curves and the corresponding AUC values were calculated to quantify intestinal amino acid absorption.

### Body composition analysis

Body fat content was assessed before pregnancy and at late gestation using a QMR body composition analyzer (QMR06-090H, Niumag Corporation) according to the manufacturer’s instructions.

### Histological analysis

Small intestinal tissues were prepared as Swiss rolls, fixed in 4% paraformaldehyde, paraffin-embedded, sectioned, and stained with hematoxylin and eosin (H&E). Whole-slide images were acquired using a NanoZoomer S360 imaging system.

### Cell proliferation tracing and fluorescence analysis

For in vivo proliferation tracing, mice were intraperitoneally injected with BrdU (Sigma-Aldrich, B5002; 50 mg/kg) 20 h before tissue collection. Intestinal tissues were fixed in 4% paraformaldehyde, paraffin-embedded, and sectioned. Tissue sections were subjected to antigen retrieval, acid denaturation, and immunostaining with anti-BrdU antibody (Abcam, ab152095), followed by fluorescent secondary antibody staining. Nuclei were counterstained with DAPI. BrdU fluorescence images were acquired using a Pannoramic SCAN II imaging system.

For organoid proliferation assays, organoids were incubated with 10 μM EdU for 8 hours before collection. Organoids were recovered from Matrigel, fixed with 4% paraformaldehyde, permeabilized, and processed using a BeyoClick™ EdU-555 Cell Proliferation Kit according to the manufacturer’s instructions. Fluorescence images were acquired using an Olympus FV4000 imaging system.

### Transmission electron microscopy

Jejunal tissues were post-fixed in osmium tetroxide, dehydrated through a graded ethanol series, embedded in resin, and sectioned into ultrathin sections (70 nm). Sections were stained with uranyl acetate and lead citrate prior to imaging using a HITACHI HT7800 transmission electron microscope.

### Villi and microvilli quantification

For histological analyses, 20–30 well-oriented intact villi were randomly selected from each sample. Villus height and mid-villus width were quantified using FIJI software. For epithelial cell area analysis, villus epithelial regions excluding the crypts and lamina propria were segmented in QuPath. Nuclei were identified using QuPath’s positive cell detection algorithm based on hematoxylin staining, and epithelial cell cross-sectional areas were estimated using the nuclear localization and cell segmentation workflow.

For microvilli analyses, transmission electron microscopy images were analyzed using the FIJI software. Microvillus length was measured from 20 randomly selected microvilli from each of 10 randomly selected fields per sample (200 microvilli per sample). Microvillus density was determined by measuring the width occupied by 15 consecutive microvilli across 10 randomly selected fields and was expressed as the number of microvilli per micrometer.

For villus height, mid-villus width, epithelial cell area, microvillus length, and microvillus density, multiple measurements were obtained per mouse and averaged to yield a single value for each mouse. Each mouse was considered an independent biological replicate, whereas individual structures and image fields were not treated as independent replicates.

### Flow cytometry

Lamina propria mononuclear cells were isolated from the small intestine after removal of Peyer’s patches and mesenteric fat. Tissues were sequentially dissociated in HBSS containing HEPES, DTT, FBS, and EDTA at 37°C with gentle shaking to remove epithelial components, followed by enzymatic digestion in RPMI 1640 supplemented with collagenase VIII, DNase I, and dispase II at 37°C. Digested tissues were filtered through 40-µm cell strainers, and immune cells were enriched by 40%/70% Percoll density-gradient centrifugation. Cells at the interphase were collected, washed, and used for downstream flow cytometric analyses.

For scRNA-seq experiments, live CD45^+^ immune cells were isolated by fluorescence-activated cell sorting (FACS). Cells were stained with Fixable Viability Stain 780 to exclude dead cells and labeled with BV510-conjugated anti-mouse CD45 antibody. Live CD45^+^ cells were sorted using a BD FACSAria™ III cell sorter (BD Biosciences) and collected for subsequent library preparation.

For Treg-cell analysis, cells were stained with Fixable Viability Stain 780, followed by surface staining with antibodies against CD45, CD4, and CD25. Cells were then fixed and permeabilized using the Foxp3/transcription factor staining buffer set according to the manufacturer’s instructions and stained intracellularly with anti-mouse Foxp3 antibody. Treg cells were defined as live CD45^+^CD4^+^CD25^+^Foxp3^+^ cells. Samples were acquired using a BD FACSVerse flow cytometer (BD Biosciences) and analyzed with FlowJo software (version X.0.7).

### ELISA assay

IL-10 concentrations in intestinal tissue lysates and cell culture supernatants were measured using commercially available Mouse IL-10 ELISA kits (Elabscience, E-EL-M0046; R&D Systems, M1000B, respectively) according to the manufacturers’ protocols. IL-10 concentrations in intestinal tissue lysates were normalized to total protein concentration.

### Dual-Luciferase reporter assays

HEK293T cells were seeded in 24-well plates and cotransfected with wild-type or mutant *Il10* promoter luciferase reporter plasmids (1 µg), *Esr1* expression plasmid or empty vector (1 µg), and Renilla luciferase control (pRL-TK, 0.1 µg) using Lipofectamine 2000. At 48 h post-transfection, luciferase activities were measured using the Dual-Luciferase Reporter Assay System (JinKaiRui, JKR23008), with firefly luciferase normalized to Renilla luciferase.

### EMSA

A biotin-labeled oligonucleotide probe (5’-ATG CAT GCA TGC ATG CAT GCA TGC-3’) was incubated with bacterial lysates expressing *Il10* at room temperature for 30 minutes. Complexes were resolved on a non-denaturing 6% polyacrylamide gel in 0.5× TBE buffer at 60 V and 4 °C for 1 hour. DNA was transferred to a Biodyne^®^ B nylon membrane (Pall Corporation) and detected using a chemiluminescent EMSA kit, visualized via ChemiDoc XRS system (Bio-Rad Laboratories, UAS).

### RNA isolation and qRT-PCR

Total RNA was extracted using a TransZol Up Plus RNA Kit (TransGen Biotech, ER501-01-V2). First-strand cDNA synthesis was performed using the TransScript^®^ All-in-One First-Strand cDNA Synthesis SuperMix (TransGen Biotech, AT341-03). Quantitative PCR was carried out using PerfectStart Green qPCR SuperMix (TransGen Biotech, AQ602-01) on a QuantStudio™ 5 (Thermo Fisher) real-time PCR system. Relative gene expression was normalized to *β-Actin* and calculated using the ΔΔCt method. All PCR primers are listed in the Figure 4-supplementary table 2.

### Bulk RNA sequencing and data analysis

Total RNA was extracted using TRIzol reagent (Invitrogen, 15596026). RNA integrity was assessed using an Agilent 2100 Bioanalyzer, and samples with an RNA integrity number (RIN) > 7 were used for library preparation. Poly(A)+ RNA was enriched with Dynabeads Oligo(dT)25 and fragmented prior to cDNA synthesis. Sequencing libraries were prepared using standard Illumina-compatible protocols and sequenced on an Illumina NovaSeq 6000 platform (PE150).

Quality control of raw sequencing data was performed using trim-galore(v-0.6.10). Transcript abundance was quantified with kallisto (v0.48.0) using the murine GRCm39 and porcine Sscrofa11.1 reference genomes with Ensembl v110 annotations. Transcript-level abundance estimates were aggregated to gene-level TPMs using the tximport R package. (v1.30.0). Protein-coding genes with TPM > 0.1 in at least 50% of samples in each condition were retained for downstream analyses. Differential expression analysis was performed using DESeq2 (v1.42.1) (***Love, Huber, & Anders, 2014***) with Wald statistics, and adjusted *P* values were calculated using Independent Hypothesis Weighting (IHW). Principal component analysis (PCA) was performed in R using the FactoMineR and factoextra packages. Enrichment of differentially expressed genes was performed using Metascape (https://pubmed.ncbi.nlm.nih.gov/30944313/) (***Zhou et al., 2019***).

### Single-cell RNA sequencing and data analysis Library preparation

CD45⁺ immune cells isolated from the intestinal lamina propria were sequentially filtered through 70-μm and 30-μm strainers, depleted of dead cells, and resuspended in PBS containing 0.04% bovine serum albumin at a final concentration of 700–1200 cells/μL. Single-cell 3′ RNA-seq libraries were generated using the MobiCube High-Throughput Single-Cell 3′ RNA Sequencing Kit v2.1 (MobiDrop) according to the manufacturer’s instructions.

### Data processing, clustering, and cell type annotation

Raw sequencing reads were aligned to the GRCm39 reference genome using MobiVision (v3.2). Ambient RNA contamination and doublets were removed using SoupX (***Young & Behjati, 2020***) and scDblFinder (***Germain, Lun, Garcia Meixide, Macnair, & Robinson, 2021***), respectively. Cells were retained if they met the following quality-control criteria: more than 350 detected genes, >1000 unique molecular identifiers (UMIs), and a mitochondrial transcript percentage below the median plus five median absolute deviations. Genes detected in fewer than 20 cells were excluded. After filtering, 65,098 high-quality cells were retained for downstream analyses. Integrated analyses were performed using Seurat (v5.1.0) (***Hao et al., 2024***). Data were normalized with NormalizeData and scaled after regressing mitochondrial transcript content and cell-cycle scores. Batch effects were corrected with Harmony using the top 2,000 highly variable genes. Dimensionality reduction, shared nearest-neighbor graph-based clustering, and UMAP visualization were performed using standard Seurat workflows. Cell identities were assigned based on canonical lineage markers. Differentially expressed genes were identified using the Wilcoxon rank-sum test with thresholds of avg_logFC > 0.25, adjusted *p* < 0.05, and min.pct > 0.20.

For T cell subclustering, integrated analyses were performed at a clustering resolution of 0.5. After quality control, a total of 62,664 high-quality cells were retained for downstream analysis. Pseudobulk expression profiles were generated using AggregateExpression, and differential expression was assessed using DESeq2 with adjusted *P*-values corrected *via* Independent Hypothesis Weighting (IHW) and log₂ fold changes stabilized using the lfcShrink function.

### Differential abundance analysis

Differential abundance analysis was conducted using miloR (v1.10.0) (***Dann et al., 2022***), a k-nearest neighbor graph-based framework. To ensure balanced comparisons between any two groups, conditions were down-sampled to equal cell numbers. A SingleCellExperiment object was created from the count matrix. Neighborhood graphs were constructed using buildGraph (k = 30, d = 50) and makeNhoods (prop = 0.2, k = 30, d = 50), yielding 6,670 representative neighborhoods.

### Cell type prioritization using Augur

Cell-type responsiveness to pregnancy was evaluated using Augur (***Skinnider et al., 2021***) at the major cell lineage level. Before analysis, up to 500 cells were randomly sampled without replacement from each condition–sample–cell subtype combination, with all cells retained in combinations containing fewer than 500 cells. Input features were restricted to the highly variable genes identified in the original Seurat object. Augur scores quantified the ability of transcriptomic profiles to distinguish pregnant (Preg) from virgin (Virg) conditions within each major cell lineage. Lineages were ranked by the area under the receiver operating characteristic curve (AUC), with higher scores indicating greater transcriptional separability between conditions.

### T cell subset proportion analysis

To evaluate alterations in T cell composition between Virg and Preg conditions, single-cell RNA sequencing data were analyzed based on annotated T cell subsets. Only T cell subsets with clearly defined cell-state annotations were included in the differential abundance analysis. The proportion of each T cell population was calculated for each biological replicate by dividing the number of cells assigned to that subset by the total number of analyzed T cells within the same replicate. Differential abundance between conditions was assessed using a quasibinomial generalized linear model, with the counts of cells belonging to each T cell subset and the remaining T cells within each replicate used as the response. The quasibinomial model was used to account for overdispersion in cell proportion data. The magnitude of changes between conditions was expressed as the log2 fold change of mean cell proportions. Statistical significance was assessed using model-derived P values, which were adjusted for multiple testing using the Benjamini– Hochberg false discovery rate (FDR) method. T cell subsets with an FDR < 0.05 were considered significantly differentially abundant between conditions.

### Transcriptional noise analysis

We applied a previously described approach to quantify transcriptional noise across cell types (***Angelidis et al., 2019***). Transcriptional noise was quantified for each cell type by analyzing a minimum of 10 cells from both Virg and Preg conditions. To standardize for differences in total UMI counts, all cells were down sampled to ensure equal library sizes. Additionally, to account for variations in cell-type frequency, scRNA-seq data were down sampled to 300 cells per cell type in each group; cell types with fewer than 300 cells included all available cells. Next, genes were categorized into ten blocks based on their average expression levels, and genes with the lowest coefficient of variation (CV) in each block (the lowest 10% CV) were selected. Euclidean distance–based scores were calculated for each cell type and condition using the expression profiles of the selected low-CV genes. These scores were used to quantify transcriptional variability and served as a proxy for transcriptional noise.

### Signature score definition

Gene set activity scores were calculated with AUCell (v1.24.0). Additional scores for IL-10– associated suppressive and *Il10* expression were derived using curated gene sets from the Gene Ontology databases.

### Cross-species integration of single-cell transcriptomics and pregnancy-related GWAS

To investigate the genetic relevance of intestinal cellular programs to pregnancy-related traits, mouse intestinal single-cell RNA sequencing (scRNA-seq) data were integrated with human genome-wide association study (GWAS) summary statistics using the scPagwas framework (***Ma et al., 2023***). Twenty-seven pregnancy-associated traits from publicly available GWAS datasets were selected for analysis (Table S1). Before integration, mouse genes were mapped to their corresponding human orthologs. A reduced Reactome pathway collection comprising 195 gene sets (reduce_genes.by.reactome.pathway) was used as the pathway input. The scPagwas framework integrates pathway activity profiles derived from single-cell transcriptomic data with GWAS summary statistics through polygenic regression to prioritize trait-associated genes and cellular populations. Trait–cell type associations were assessed across annotated intestinal cell populations, and trait relevance at the single-cell level was evaluated using the TRS output of scPagwas.

For gene-level association analysis within regulatory T (Treg) cells, Treg cells were extracted from the intestinal scRNA-seq dataset, and the scPagwas workflow was rerun separately on this subset for each pregnancy-related trait using the same Reactome pathway collection. Pathway activity profiles and cell-level gPAS scores were recalculated using the Treg subset. For each gene, a Pearson correlation coefficient (PCC) was calculated across Treg cells between its normalized expression and the corresponding gPAS scores. These PCCs were used to prioritize candidate genes whose expression was associated with pregnancy-related genetic signals within the Treg population.

### Blinding

To avoid analytical bias, investigators were blinded to group allocation for specific assays, including omics data analysis, flow cytometry, intestinal histomorphometry, nutrient absorption assays, EMSA, and dual-luciferase reporter assays. Conversely, blinding was not performed for the remaining validation experiments.

### Quantification and Statistical Analysis

Statistical analyses were performed using GraphPad Prism 9. Data are presented as means ± SEM unless otherwise indicated. Comparisons between two groups were analyzed using two-tailed unpaired Student’s *t*-tests. Comparisons among multiple groups were performed using one-way ANOVA followed by Tukey’s multiple-comparison test. Differences were considered statistically significant at *p* < 0.05.

## Key Resources Table

| REAGENT OR RESOURCE | SOURCE | IDENTIFIER |
|---|---|---|
| Antibodies |  |  |
| Anti- BrdU antibody | Abcam | Cat#ab152095; RRID:AB_2813902 |
| Alexa Fluor- 594 conjugated donkey anti- rabbit secondary antibody | Invitrogen | Cat#A-21207; RRID: AB_141637 |
| Fixable Viability Stain 780 | BD Biosciences | Cat#565388; RRID:AB_2869673 |
| BV510 rat anti-mouse CD45 | BD Biosciences | Cat#563891; RRID:AB_2734134 |
| BB700 rat anti-mouse CD4 | BD Biosciences | Cat#566407; RRID:AB_2744427 |
| BV421 rat anti-mouse CD25 | BD Biosciences | Cat#564571; RRID:AB_2738849 |
| Anti-mouse Foxp3 APC | eBioscience<sup>TM</sup> | Cat#17-5773-82; RRID:AB_469457 |
| Purified NA/LE Hamster Anti-Mouse CD3e | BD Biosciences | Cat#553057; RRID:AB_394590 |
| Purified NA/LE Hamster Anti-Mouse CD28 | BD Biosciences | Cat#553294; RRID:AB_394763 |
| Aanti-Mouse IL- 10R/CD210 Antibody (1B1.3A) | MCE | Cat#HY- P990228 |
| Chemicals, peptides, and recombinant proteins |  |  |
| Diphtheria Toxin | Sigma-Aldrich | Cat#D0564 |
| Estradiol | Selleck | Cat#S1709 |
| Progesterone | Selleck | Cat#S1705 |
| Glucose | Sigma-Aldrich | Cat#G7021 |
| BrdU | Sigma-Aldrich | Cat#B5002 |
| DAPI | Solarbio | Cat#C0060 |
| DPBS | Gibco | Cat#C14190500BT |
| Matrigel | BioGenous | Cat#M315077 |
| Recombinant mouse IL- 10 protein (active) | Abcam | Cat#ab281800 |
| Immunostaining blocking buffer | Beyotime | Cat#P0260 |
| Permeabilization solution | Beyotime | Cat#P0097 |
| Antifade Mounting Medium containing DAPI | Beyotime | Cat#P0131 |
| Hanks' Balanced Salt Solution (HBSS) | Absin | Cat#abs9257 |
| HEPES | Yuanye Bio | Cat#R27174 |
| DTT | Yuanye Bio | Cat#R21089 |
| Fetal Bovine Serum (FBS) | Gibco | Cat#10091-148 |
| EDTA | Yuanye Bio | Cat#R21090 |
| RPMI 1640 | Gibco | Cat#22400-089 |
| collagenase VIII | Sigma-Aldrich | Cat#C2139 |
| DNase I | Worthington | Cat#LS002140 |
| Dispase II | Sigma-Aldrich | Cat#D4693 |
| Percoll | Cytiva | Cat#17089109 |
| Stain Buffer | BD Pharmingen | Cat#554656 |
| Recombinant Mouse IL-2 Protein | R&D Systems | Cat#402-ML-020 |
| Recombinant Mouse TGF- β Protein | R&D Systems | Cat#240-B-002 |
| ICI- 182780 | Selleck | Cat#S1191 |
| Trizol | Invitrogen | Cat#15596026 |
| Dynabeads Oligo(dT)25 | Invitrogen | Cat#61005 |
| Magnesium RNA fragmentation module | NEB | Cat#E6150S |
| SuperScript II reverse transcriptase | Invitrogen | Cat#18064071 |
| E. coli DNA polymerase I | NEB | Cat#M0209 |
| RNase H | Thermo Scientific | Cat#EN0201 |
| AMPure XP beads | Beckman Coulter | Cat#A63880 |
| Red Blood Cell Lysis Solution | Miltenyi Biotec | Cat#130- 094- 183 |
| Dead Cell Removal Kit | Miltenyi Biotec | Cat#130- 090- 101 |
| Lipofectamine™ 2000 Transfection Reagent | Invitrogen | Cat#11668019 |
| Lipofectamine™ 3000 Transfection Reagent | Invitrogen | Cat#L3000015 |
| HRP-conjugated Streptavidin | Proteintech | Cat#SA00001-0 |
| EMSA/Gel-Shift Binding Buffer (5X) | Beyotime | Cat#GS005 |
| EMSA/Gel-Shift Loading Buffer (10X) | Beyotime | Cat#GS006 |
| Critical commercial assays |  |  |
| Mouse Intestinal Organoid Kit (Serum-free) | BioGenous | Cat#K2001-MI |
| BeyoClick™ EdU-555 Cell Proliferation Kit | Beyotime | Cat#C0075S |
| Foxp3/Transcription Factor Staining Buffer Set | eBioscience™ | Cat#00-5523-00 |
| Naive CD4<sup>+</sup> T Cell Isolation Kit | Miltenyi Biotec | Cat#130-104-453 |
| Mouse Il-10 (Interleukin 10) elisa kit | Elabscience | Cat#E-EL-M0046 |
| Mouse Il-10 quantikine elisa kit | R&D Systems | Cat#M1000B |
| TransZol Up Plus RNA Kit | TransGen Biotech | Cat#ER501-01-V2 |
| TransScript All--in--One First-Strand cDNA Synthesis SuperMix for qPCR | TransGen Biotech | Cat#AT341-03 |
| PerfectStart Green qPCR SuperMix | TransGen Biotech | Cat#AQ602- 01 |
| MobiCube high- throughput single- cell 3′ RNA sequencing kit v2.1 | MobiDrop | Cat#S050200301 |
| Universal DNA Purification Kit | TIANGEN | Cat#DP214 |
| Double-Luciferase Reporter Assay Kit | JinKaiRui | Cat#JKR23008 |
| Deposited data |  |  |
| Porcine small intestine bulk RNA sequencing | This paper | GSA: CRA047091 |
| Murine small intestine bulk RNA sequencing | This paper | GSA: CRA047065 |
| Single-cell sequencing of FACS sorted small intestinal lamina propria CD45+ Cells | This paper | GSA: CRA047125 |
| Experimental models: Organisms/strains |  |  |
| Mouse: C57BL/6J | Vital River | N/A |
| Mouse: *Foxp3<sup>DTR</sup>* | Jackson Laboratory | Cat#16958 |
| Mouse: *Esr1<sup>fl/fl</sup>* | Jackson Laboratory | Cat#032173 |
| Mouse: *Foxp3-Cre* | Cyagen | Cat#C001467 |
| Software and algorithms |  |  |
| GraphPad Prism 10.6.0 | GraphPad Software | RRID:SCR_002798 |
| Adobe Illustrator | Adobe | RRID: SCR_010279 |
| FACSDiva v9.0.1 | BD Biosciences | RRID:SCR_001456 |
| FlowJo v10.0.7 | BD Biosciences | RRID:SCR_008520 |
| Fiji (imageJ) v2.3.0 | National Institutes of Health | RRID:SCR_002285 |
| QuPath v0.7.0 | https://qupath.github.io | RRID:SCR_018257 |
| Analyst®TF | SCIEX | RRID:SCR_015785 |
| kallisto v0.48.0 | https://pachterlab.github.io/kallisto | RRID:SCR_016582 |
| tximport v1.30.0 | https://bioconductor.org/packages/release/bioc/html/tximport.html | RRID:SCR_016752 |
| DESeq2 v1.42.1 | https://bioconductor.org/packages/release/bioc/html/DESeq2.html | RRID:SCR_015687 |
| FactoMineR v2.11 | https:/www.R-project.org | RRID:SCR_014602 |
| factoextra v1.0.7 | https://github.com/kassambara/factoextra | RRID:SCR_016692 |
| MobiVision v3.2 | https://www.mobidrop.com | N/A |
| SoupX v1.6.2 | https://github.com/constantAmateur/SoupX | RRID:SCR_019193 |
| scDblFinder v1.16.0 | https://github.com/plger/scDblFinder | RRID:SCR_022700 |
| Seurat v5.1.0 | https://satjalab.org/seurat/ | RRID:SCR_016341 |
| miloR v1.10.0 | https://github.com/MarioniLab/miloR | RRID:SCR_025630 |
| AUCell v1.24.0 | https://github.com/aertslab/AUCell | RRID:SCR_021327 |
| scPagwas | https://github.com/sulab-wmu/scPagwas | N/A |
| Other |  |  |
| NanoZoomer S360 | Hamamatsu | N/A |
| Pannoramic SCAN II | 3DHISTECH | N/A |
| HT7800 | HITACHI | N/A |
| FV4000 | Olympus | N/A |
| gentleMACS Dissociator | Miltenyi Biotec | N/A |
| FACSAria™III | BD Biosciences | N/A |
| FACSVerse™ | BD Biosciences | N/A |
| Countstar Rigel | Alit Biotech | N/A |
| NovaSeq 6000 | Illumina | N/A |
| Bioanalyzer 2100 | Agilent | N/A |
| NanoDrop ND-1000 | Thermo Fisher | N/A |
| 1290 II UPLC | Agilent | N/A |
| 5600+ quadrupole-TOF | Sciex | N/A |
| QuantStudio™ 5 | Thermo Fisher | N/A |
| ChemiDoc XRS system | Bio-Rad Laboratories | N/A |
| Chemiluminescence immunoassay analyzer BK-L96C | Kinghawk | N/A |
| QMR06- 090H- PRO low- field NMR small- animal body composition analyzer | Niumag | N/A |

## Author contributions

L.H.: Conceptualization, Methodology, Investigation, Validation, Writing – original draft, Writing – review & editing, Funding acquisition, Supervision. M.L.: Conceptualization, Methodology, Funding acquisition, Writing – review & editing. D.W.: Conceptualization, Methodology, Funding acquisition, Writing – review & editing. L.Liu.: Methodology, Investigation, Data curation, Validation, Visualization, Writing – original draft, Writing – review & editing. X.Jin.: Data curation, Validation, Visualization, Writing – review & editing. J.Z.: Investigation, Data curation. Y.W.: Investigation. S.T.: Investigation. L.Lu.: Investigation. Y.Z.: Investigation. C.J.: Investigation. B.F.: Investigation. L.C.: Investigation. S.X.: Investigation. Y.L.: Investigation. L.J.: Investigation. X.Jiang.: Project administration.

## Data availability

All data and code needed to evaluate and reproduce the conclusions in the paper are present in the paper and/or the Supplementary Materials. The high-throughput sequencing datasets generated in this study have been deposited in the Genome Sequence Archive (GSA) (https://ngdc.cncb.ac.cn/gsa/) of the China National Center for Bioinformation National Genomics Data Center (CNCB-NGDC) under accession numbers CRA047065 (murine RNA-seq data), CRA047091 (porcine RNA-seq data) and CRA047125 (murine scRNA-seq data). All other materials are available from the corresponding author upon reasonable request.

## Additional files

Figure 1-supplementary figure 1

Figure 2-supplementary figure 2

Figure 3-supplementary figure 3

Figure 3-supplementary figure 4

Figure 4-supplementary figure 5

Figure 5-supplementary figure 6

Figure 6-supplementary table 1

Figure 4-supplementary table 2

MDAR checklist

